# An Expectation and Maximization Algorithm for Multivariate Genome-wide Association Studies (EMmvGWAS)

**DOI:** 10.64898/2025.12.02.691906

**Authors:** Chin-Sheng Teng, Xuesong Wang, Cheng Liu, Qishan Wang, Yanru Cui, Shizhong Xu

## Abstract

Genome-wide association studies (GWAS) commonly focus on one quantitative trait at a time, even when multiple traits are collected. However, joint analysis of multiple traits can increase the power to detect genetic associations by leveraging trait correlations. Multivariate GWAS is particularly important for identifying pleiotropic effects and uncovering shared genetic architecture underlying complex traits. Despite its great potential, multivariate GWAS methods face substantial computational challenges due to the high dimensionality of polygenic covariance structures and the cost of scanning genome-wide markers. We present EMmvGWAS, an efficient computational framework implemented as an R, which dramatically reduces computation time for multivariate GWAS involving a moderate number of traits. The algorithm estimates the genetic and environmental covariance matrices from a pure polygenic model using an Expectation-Maximization (EM) algorithm. To improve computational efficiency during genome-wide scanning, we use a semi-exact method that fixes the estimated covariance matrices and treats their ratio, defined as the genetic covariance matrix multiplied by the inverse of the environmental covariance matrix, as a known constant across all markers. This approach enables closed-form solutions for marker effects at each locus, dramatically reducing the computation time without compromising statistical power. We demonstrate the scalability and performance of our method through both simulation studies and real dataset analyses from rice, mouse and human populations. The method is implemented in R and is designed to support multivariate GWAS involving a moderate number of traits, while also accommodating univariate analyses as a special case. The R package is available at https://github.com/Jason-Teng/EMmvGWAS.

## 1. Background

Genome-wide association studies (GWAS) have significantly advanced our understanding of the genetic architecture underlying complex traits by identifying genetic variants associated with quantitative traits (Visscher et al., 2017). Traditionally, GWAS are often conducted on individual traits independently, even though multiple correlated phenotypes are always measured in genetic studies. Analyzing traits individually overlooks valuable trait correlations, thus potentially reducing the statistical power and increasing the type 1 error rate. Multivariate GWAS leverages these correlations by using multivariate linear mixed models, improving power and reducing false positive associations in identifying pleiotropic genetic variants, which affect multiple traits simultaneously (Korte et al., 2012; Porter & O’Reilly, 2017; Turley et al., 2018; Zhou & Stephens, 2014). Such methods are crucial for uncovering shared genetic bases among related traits, providing deeper insights into biological pathways and mechanisms underlying complex diseases and phenotypes (Hackinger & Zeggini, 2017).

Despite their clear advantages, multivariate GWAS methods confront significant computational hurdles, primarily due to the complexity of modeling high-dimensional covariance structures and the computational cost associated with scanning genome-wide markers. Estimating genetic and environmental covariance matrices between traits becomes increasingly demanding as the number of traits grows, often requiring iterative optimization over large parameter spaces. At the same time, genome-wide marker scanning involves evaluating potentially millions of SNPs across all traits, which remains computationally intensive. These two challenges compound in high-dimensional settings, making many existing multivariate GWAS tools impractical for real-world applications involving many traits. Although R is widely used in statistical genetics, there is a lack of efficient R packages for multivariate GWAS, especially for analyses involving many traits.

To address these challenges, we developed EMmvGWAS, a general statistical framework that supports multivariate GWAS with improved computational efficiency. While suitable for small trait sets, EMmvGWAS offers substantial speed improvements as the number of traits grows, making it especially effective for studies involving 6 to 10 traits or more. For variance component estimation, we applied an approach similar to previous methods that utilize eigen decomposition of the genetic relationship matrix (or kinship matrix) and linear transformation to achieve uncorrelated observations. However, we simplify the model into a weighted linear mixed model with a random intercept, allowing for more efficient estimation. Variance components are estimated using an expectation-maximization (EM) algorithm under a pure polygenic model.

For marker scanning, we developed a semi-exact method using an innovative linear transformation framework. Traditional approximate approaches use variance components estimated under the null model, offering faster computation but at the cost of reduced statistical power and lacking a natural extension to multivariate settings (Kang et al., 2010; Zhang et al., 2010). In contrast, exact methods such as GEMMA maintain statistical rigor but become computationally burdensome as the number of traits increases (Zhou & Stephens, 2014). Our EMmvGWAS algorithm addresses these limitations by using a semi-exact method that fixes the ratio of variance components while explicitly re-estimating the residual covariance matrix at each locus. This enables a closed-form Wald test for each marker, eliminating the need for iterative optimization during marker testing and substantially reduces time complexity, achieving a practical balance between computational efficiency and statistical power.

To promote accessibility and reproducibility, we implemented the method in R, a widely used platform in statistical genetics. Our package supports moderate-dimensional multivariate GWAS with flexible input for phenotypes, covariates, and kinship matrices, enabling scalable multivariate analyses and deeper insight into the genetic architecture of complex traits.

## 2. Methods

### 2.1. Multivariate linear mixed model

The multivariate linear mixed model for *p* traits measured on *n* individuals is defined as:

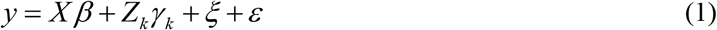

where the phenotype vector is stacked by individuals, 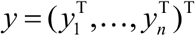, and *y* = ( *y*_*j*1_, …, *y*_*jp*_)^T^ is the *p* ×1 vector of trait values for individual *j, j* = 1, …, *n* . To test for associations, the model considers one marker at a time. For the *k* th marker, the design matrix is *Z*_*k*_ = *z*_*k*_ ⊗ *I* _*p*_, where ⊗ denotes the Kronecker product, *z*_*k*_ is the *n* -dimensional vector of genotypes for that specific marker, and *γ*_*k*_ = (*γ*_*k*1_, …,*γ*_*kp*_)^T^ is the vector of its trait-specific effects. The fixed effects design matrix *X* = 1_*n*_ ⊗ *I* _*p*_ contains trait-specific intercepts and the corresponding vector of fixed effects is *β* = (*β*_1_, …, *β*_*p*_)^T^ . The random polygenic effects and residual errors are denoted by 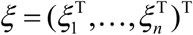 and 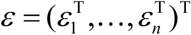 respectively, where each *ξ*_*j*_ and *ε*_*j*_ is a *p* -dimensional vector corresponding to individual *j* . We assume var(*ξ* ) = *K* ⊗∑_*ξ*_ and var(*ε* ) = *I*_*n*_ ⊗∑_*ε*_, where ∑_*ξ*_ and ∑_*ε*_ are *p* × *p* symmetric positive-definite matrices describing trait-level polygenic and residual covariances, respectively, and *K* is the known kinship matrix. Consequently, the total variance of the observed phenotypes is expressed as:

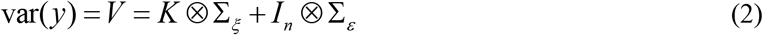

This structured covariance allows the model to capture genetic background effects and environmental noise across all *p* traits simultaneously. The inclusion of correlated individuals substantially increases computational complexity, primarily due to the necessity of repeated inversions and determinant calculations of the large-scale matrix *V* during variance component estimation. Therefore, we propose a decorrelation method to render individuals uncorrelated to simplify the inversion of covariance matrices.

### 2.2. Decorrelation of the polygenic effects

Let the eigen-decomposition of the kinship matrix be *K* = *UDU*^*T*^, where *D* = *diag*{*d*_1_,… *d*_*n*_} is the diagonal matrix holding the ordered eigenvalues and *U* is the matrix holding the eigenvectors corresponding to the eigenvalues. We define the transformation matrix as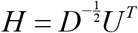 Since *y* is a stacked vector of phenotypic values across *n* individuals and *p* traits, the transformation is applied to all variables in the model using the Kronecker product *H* ⊗ *I* _*p*_ .The transformed variables are defined as *y* = (*H* ⊗ *I* _*p*_ ) *y, X* = (*H* ⊗ *I* _*p*_ ) *X, Z*_*k*_ = (*H* ⊗ *I* _*p*_ )*Z*_*k*_, *ξ* = (*H* ⊗ *I* _*p*_ )*ξ*, and *ε* = (*H* ⊗ *I* _*p*_ )*ε* . Note that we have used a pseudo code notation to simplify the derivation. Hereafter, all variables in the linear mixed model are eigen-transformed. The phenotypic variance-covariance matrix of the transformed model is given by:

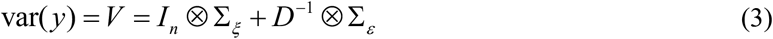

The matrix *V* becomes block diagonal with n blocks of dimension *p* × *p* . Compared with the original variance structure, the transformed covariance matrix is substantially simplified, as observations from different individuals are uncorrelated. For the *j* th individual, let *y*_*j*_ = ( *y*_*j*1_, …, *y*_*jp*_)^T^ denote the vector of transformed phenotypic values for the *p* traits. The corresponding polygenic effects and residual errors are denoted by *ξ* _*j*_ and *ε* _*j*_, respectively, where *ξ* _*j*_ and *ε* _*j*_ are *p* ×1 vectors for *j* = 1, …, *n* . The transformed model for the *j* th individual is

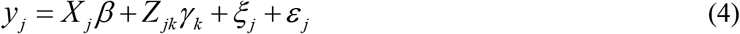

where *X* _*j*_ and *Z* _*jk*_ denote the transformed fixed-effect and marker design matrices corresponding to transformed individual *j*, respectively. The variance of *y*_*j*_ is

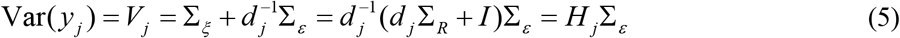

Where 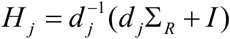, and 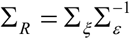 is the matrix version of variance ratio.

### 2.3 The expectation and maximization (EM) algorithm for estimating the covariance matrices

The decorrelation of polygenic effects makes all individuals independent. As a result, the log-likelihood function only involves matrices of dimension *p* × *p* for *p* traits. We estimate the covariance matrices under the null model, which includes only fixed intercepts and polygenic effects and excludes marker effects. Let *θ* = {∑_*ξ*_, ∑_*ε*_ } be the parameter set, while the *β* vector is not a vector of parameters in REML (it is a vector of parameter for ML). The restricted log likelihood function is

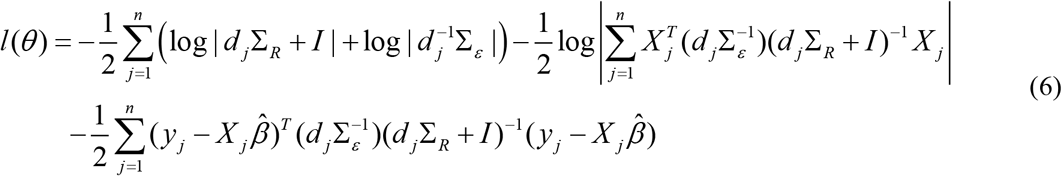

Where

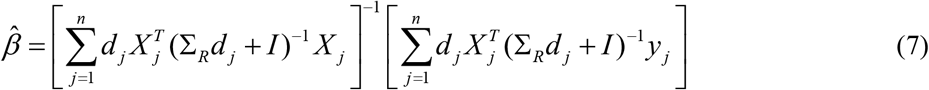

Here, 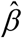 is not a parameter vector but a function of *θ* . The covariance parameters are obtained by maximizing *l*(*θ* ) . The MIXED procedure PROC MIXED in SAS is readily available to perform the variance component estimations. However, PROC MIXED becomes computationally inefficient and may fail to converge properly due to the large number of parameters. To address these issues, we adopt an Expectation-Maximization (EM) algorithm. The EM algorithm guarantees a monotonic increase in the likelihood and achieves stable convergence.

In the EM algorithm framework, the missing values in the complete-data log likelihood function are the polygenic effects, *ξ* _*j*_ for *j* = 1, …, *n*, where each *ξ* _*j*_ is an *p* ×1 vector. We first update *β* using

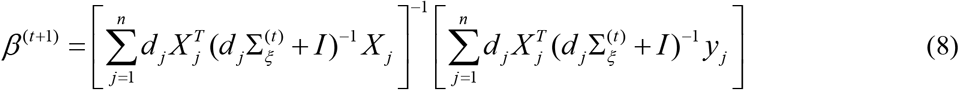

Given *ξ* _*j*_ for *j* = 1, …, *n*, the polygenic variance and covariance matrix is define as

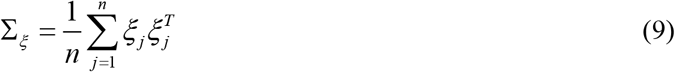

When *ξ* _*j*_ is missing, we simply replace them by the conditional expectations to update this matrix,

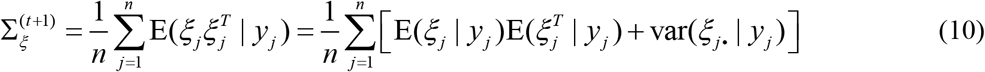

Where

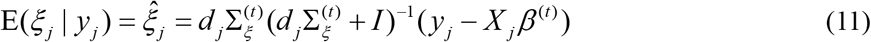

And

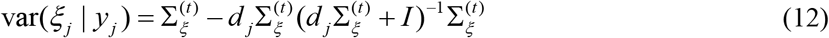

Finally, the updated residual covariance matrix is

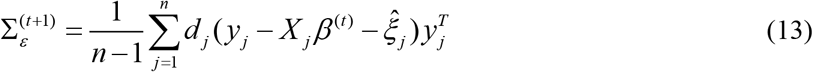

When the iterations converge, we report the two covariance matrices as the estimated parameters denoted by 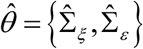 and the estimated covariance matrix ratio denoted by 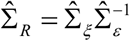. This matrix ratio will be treated as a constant and used in genome-wide scanning for associations of genome-wide markers with multiple traits.

### 2.4 Scanning the entire genome for associations

Genome-wide association analysis is carried out by testing markers one at a time. In principle, marker effects can be incorporated into the model by re-estimating all parameters for each marker, yielding an exact but computationally expensive approach. In this study, we develop an efficient strategy that fixes the variance matrix ratio estimated from the null model and scans the genome marker by marker. This strategy allows closed-form estimation of marker effects and enables re-estimation of the residual covariance matrix for each marker, thereby substantially improving computational efficiency while maintaining good statistical performance. With the variance matrix ratio fixed from the null model, genome-wide association testing is carried out marker by marker. For marker *k*, the fixed effects include both the intercepts *β* and the marker effects *γ* _*k*_, which are estimated jointly by

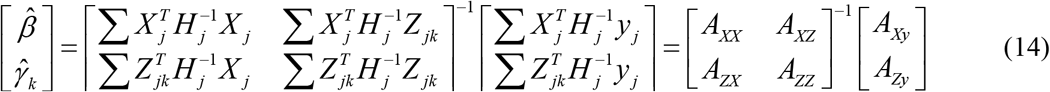

where 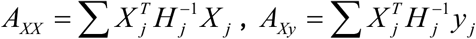, and similarly for the remaining block terms.

The intercepts *β* are nuisance parameters and can be eliminated prior to estimating *γ* _*k*_ via the annihilator matrix (Xu, 2024) so that *γ* _*k*_ are estimated explicitly. Let us also define

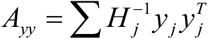. Define the following quantities

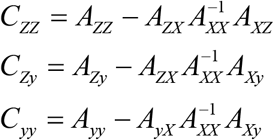

The estimated *γ* _*k*_ is

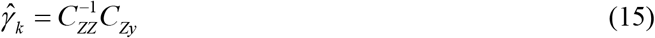

with variance

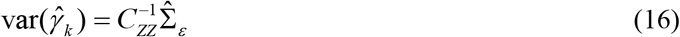

Where

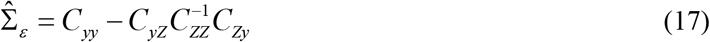

is the estimated residual covariance matrix and 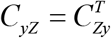. The Wald test statistic for testing *H*_0_ : *γ* _*k*_ = 0 is

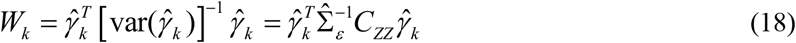

which follows approximately a Chi-square distribution with *p* degrees of freedom.

### 2.5. Dataset

We evaluated the performance and computational efficiency of the new method using three genetic association datasets from rice, human, and mouse populations. Each dataset includes genome-wide genotype data and multiple phenotypic traits, enabling a range of multivariate GWAS applications. The rice dataset consists of 1,495 hybrid rice individuals (Huang et al., 2015). For the human dataset, we analyzed a subset of 5,000 unrelated white British individuals from the UK Biobank (Sudlow et al., 2015) cohort for demonstration purposes. The mouse dataset was obtained from the CFW outbred population study (Parker et al., 2016), available from the Dryad Digital Repository (doi:10.5061/dryad.2rs41).

For the rice and mouse datasets, we applied standard quality control procedures, including removal of SNPs with minor allele frequency below 0.01 and individuals or variants with missing data. For the UK Biobank data, genotype data were filtered using PLINK2 with thresholds of 0.05 for missing genotype rate and minor allele frequency, along with linkage disequilibrium pruning (Chang et al., 2015). Detailed information is provided in Supplementary Table S1. All phenotypes were standardized prior to analysis.

### 2.6. Simulation analysis

To evaluate the type I error control and statistical power of our method, we performed null simulations and alternative simulations using real genotype data from the rice and mouse datasets. We excluded the human dataset from simulations due to its large sample size and dense marker coverage, which made simulations computationally impractical.

For null simulations, phenotypes were generated under a pure polygenic model without any marker effects. Specifically, we simulated 100 sets of phenotypes based on the real genetic relatedness matrix and the estimated genetic and environmental covariance matrices from the rice and mouse data. We randomly selected 10,000 SNPs from the real genotype data. We then computed p-values for all combinations of these SNPs and the simulated phenotypes, resulting in one million tests per dataset. For alternative simulations, we added SNP effects into the phenotypes to assess empirical power. We first selected 1,000 evenly spaced SNPs from a set of SNPs that showed no significant association with either trait (P > 0.1 in all univariate and multivariate tests). Each selected SNP was assigned an effect on the first trait to explain a fixed proportion of phenotypic variance (PVE). The effect on the second trait was scaled to explain either 20% or 40% of PVE of the second trait, with effects either in the same or opposite direction across traits. These synthetic effects were added to the original phenotypes to generate datasets with known causal SNPs. Power was calculated as the proportion of causal SNPs detected at the Bonferroni-corrected significance level with initial significance level of 0.05. We then applied both EMmvGWAS and GEMMA to these simulated datasets to compare their empirical power under various effect sizes and trait correlation scenarios.

## 3. Results

### 3.1. Computational results

To evaluate the computational performance of our method, we presented its theoretical time complexity in comparison with existing approaches. EMmvGWAS benefits from an explicit solution, resulting in a reduced overall computational complexity of *O*(*n*^3^ + *n*^2^ *p* + *t*_1_ *np*^2^ + *sn*^2^ + *snp*^2^ ), where *n* is the sample size, *p* is the number of traits, *t*_1_ is the number of iterations of the EM algorithm, and *s* is the number of markers. The terms *n*^3^ and *n*^2^*d* arise from eigen decomposition and transformation of the phenotypes. The term *t*_1_ *np*^2^ corresponds to the variance component estimation under the null model using the EM algorithm, while *s*(*n*^2^ + *np*^2^ ) reflects the cost of genome-wide marker scanning across multiple traits. In practice, the number of markers *s* is typically much larger than both *n* and *p*, making the last term *s*(*n*^2^ + *np*^2^ ) the dominant contributor to the overall complexity. The number of fixed effects per test is omitted in these expressions because its contribution is negligible and can be adjusted in the phenotypes. In contrast, the exact algorithms such as GEMMA require *O*(*n*^3^ + *n*^2^ *p* + *sn*^2^ + *st*_1_ *np*^2^ + *st*_2_*np*^2^ ), where *t*_1_ and *t*_2_ represent the number of optimization iterations needed for the EM algorithm and the Newton-Raphson algorithm, respectively. As the number of traits increases, the additional iterative steps in genome marker scanning in GEMMA lead to substantially higher computational burden.

We evaluated the computational efficiency of EMmvGWAS using real-world datasets from rice, mouse, and human, comparing it against GEMMA, the gold standard exact algorithm. For each dataset, we varied the number of traits and recorded the total runtime. As shown in Figure 1, the runtime of GEMMA increases rapidly with the number of traits. For example, GEMMA required more than three weeks (554 hours) for 10 traits in rice and nearly 200 hours for 10 traits in the human dataset. In contrast, our EMmvGWAS maintains a nearly flat runtime profile, requiring less than 5 hours for the rice and the mouse data. For the human dataset, we applied 10-core parallel computing to EMmvGWAS, which enabled analysis of up to 10 traits within 40 hours, compared to nearly 200 hours for GEMMA.

**Figure 1.**
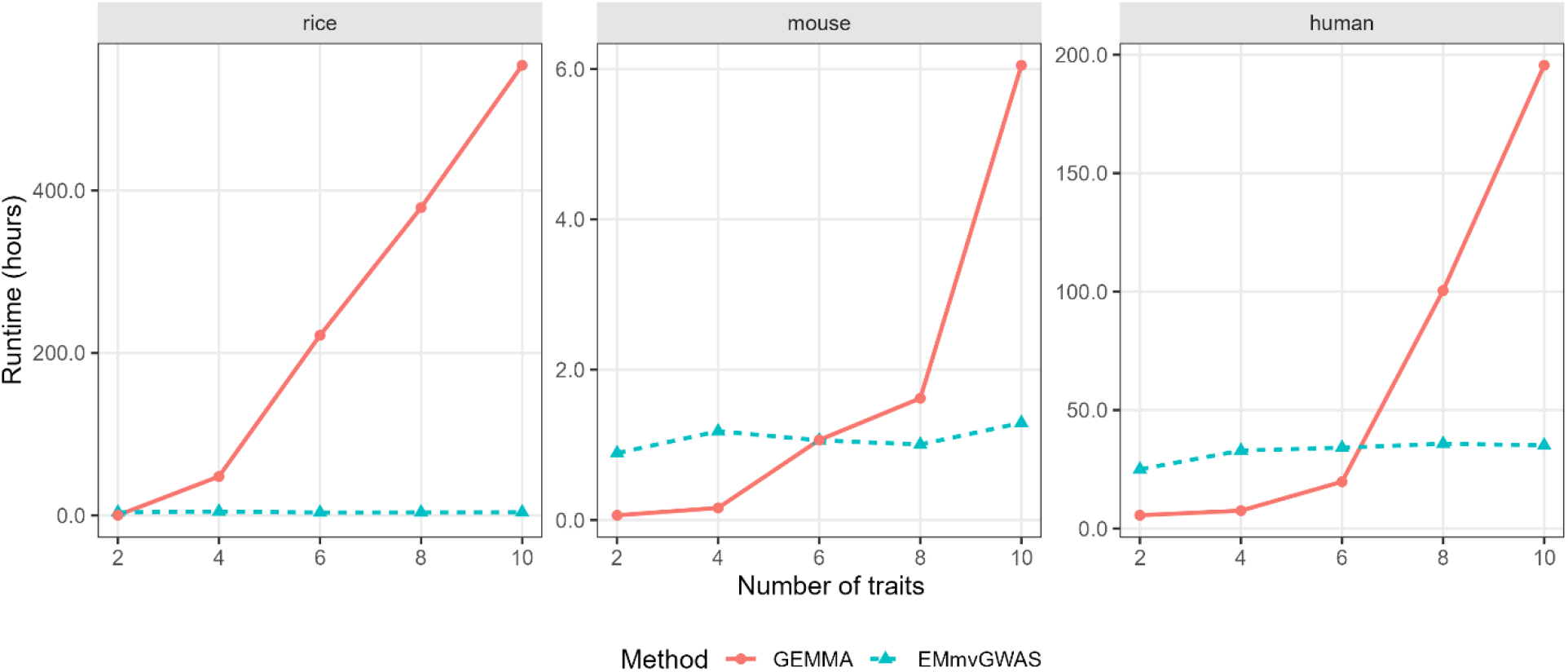
Computational runtime of EMmvGWAS and GEMMA across datasets and trait dimensions. Runtime (in hours) is plotted across the number of traits for the rice, the mouse, and the human datasets. The red solid line indicates GEMMA and the blue dashed line indicate EMmvGWAS. The rice and mouse analyses were carried out on 1 core, while the human data analysis was run with 10 cores.

### 3.2. Consistency of association results

We showed that the semi-exact method EMmvGWAS is highly consistent with the exact method implemented in GEMMA when analyzing 10 traits across the rice, the mouse, and the human datasets (Figure 2). The Pearson correlation coefficients were nearly 1 in all cases, indicating that the two approaches produced almost identical test statistics. Figure 3 displays the Manhattan plots from both methods, which show highly similar peak patterns throughout the genome, further confirming the alignment of their results.

**Figure 2.**
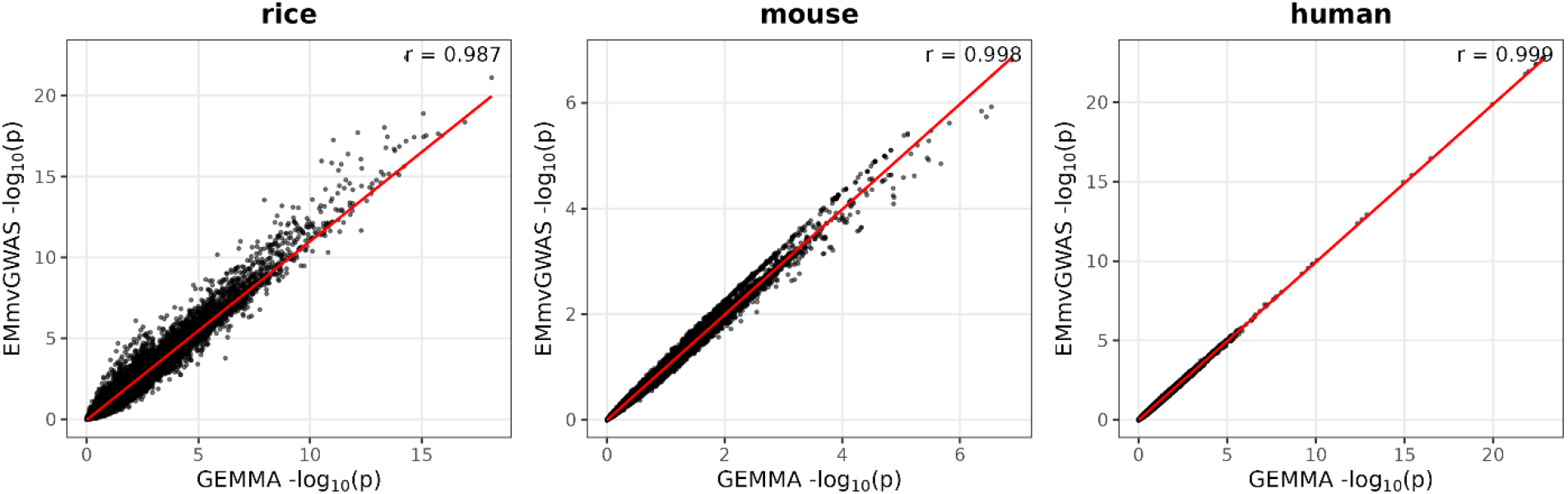
Comparison of −log10(p) values between the EMmvGWAS and GEMMA for 10 traits. Scatterplots of EMmvGWAS versus GEMMA results are shown for the rice, the mouse, and the human datasets. The red line represents the fitted regression line, and *r* denotes the Pearson correlation coefficient between methods.

**Figure 3.**
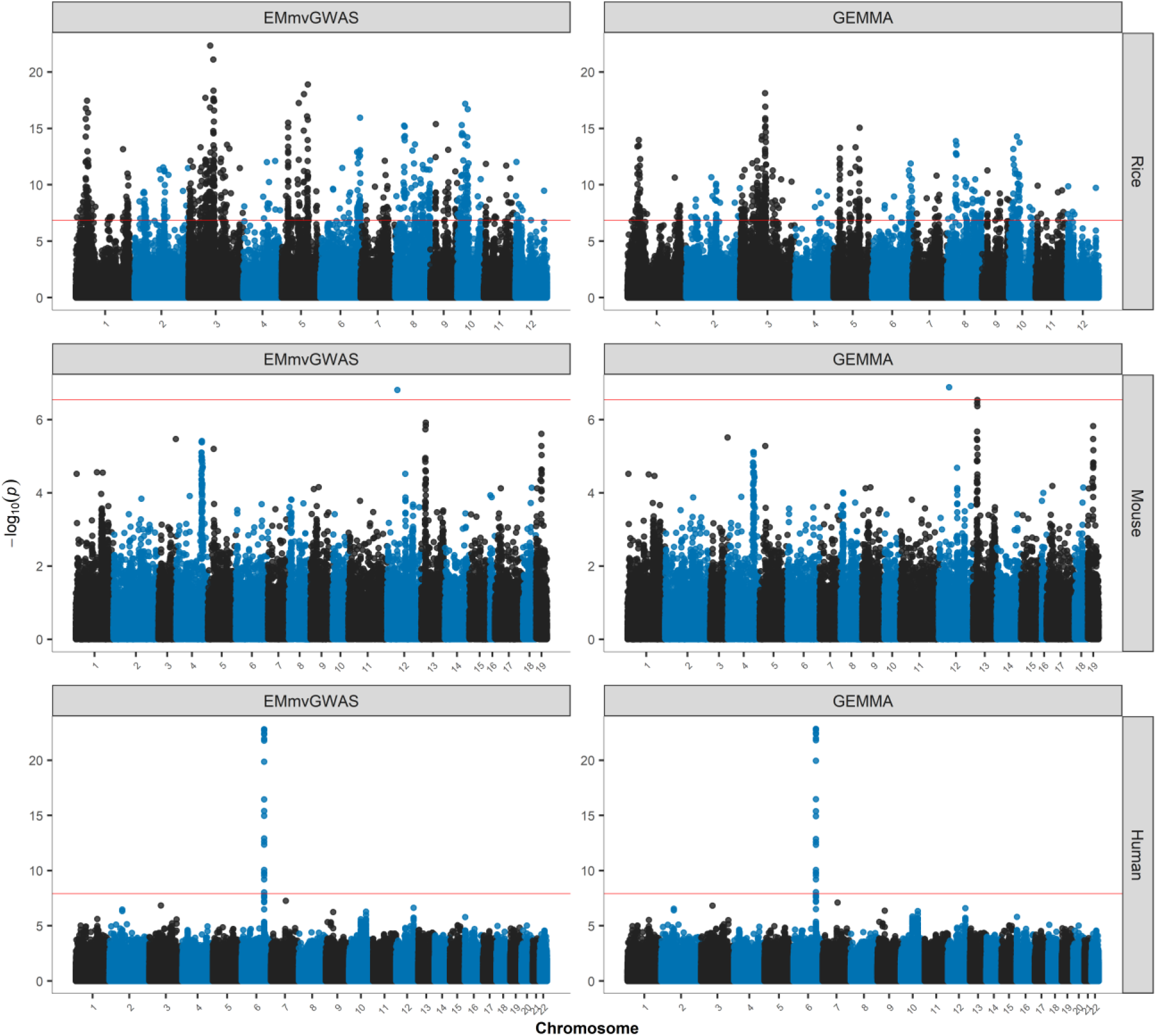
Manhattan plots for EMmvGWAS and GEMMA methods across rice, mouse, and human datasets using 10 traits. Each row shows results for one species (rice, mouse, and human), and each column corresponds to a method (EMmvGWAS on the left and GEMMA on the right). Genome-wide association −log10(p) values are plotted along the genome. The red horizontal line marks the Bonferroni-corrected significance threshold for each species.

### 3.3. Simulation study

To illustrate that our method produces P values with proper control of type 1 error and power, we conducted both null and alternative simulations based on real genotype data from the rice and mouse datasets. Under the null model, simulated phenotypes were generated without any marker effects. Quantile–quantile (QQ) plots from the null simulations in both the mouse and the rice data are shown in Figure 4. For both EMmvGWAS and GEMMA, the observed p-values closely followed the expected distribution under the null hypothesis, with only minor deviations in the extreme tail. This indicates that both approaches properly control the type 1 error across species. To evaluate statistical power, we conducted alternative simulations by injecting known genetic effects into selected SNPs with predefined proportions of variance explained. In Figure 5, GEMMA exhibited slightly higher, EMmvGWAS maintained competitive performance with minimal power loss. Importantly, EMmvGWAS achieved this with substantially reduced computational burden, demonstrating its practical advantage for multivariate GWAS applications.

**Figure 4.**
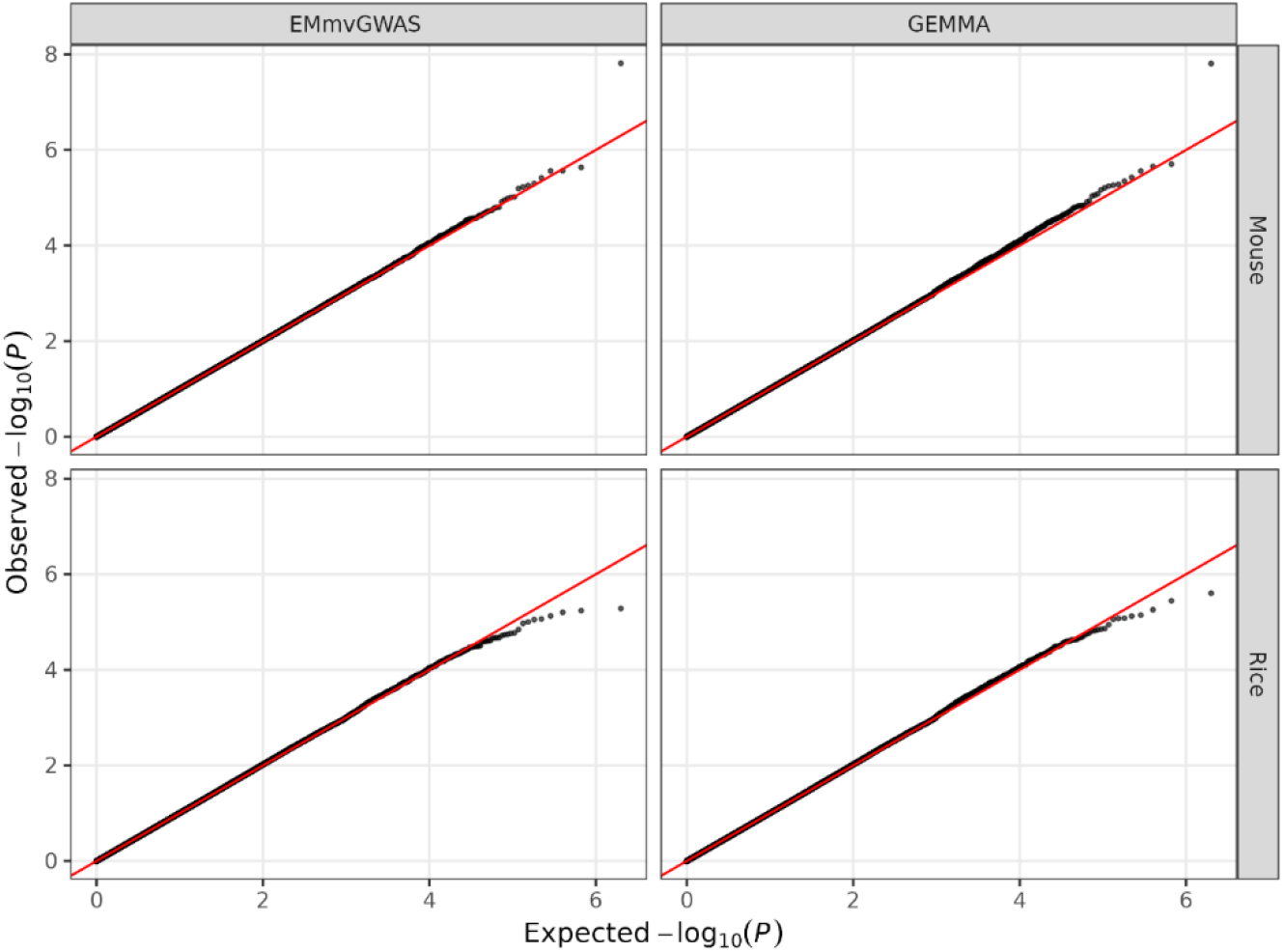
QQ plots for null simulations in the mouse and the rice data. Observed versus expected −log10(p) values are shown for EMmvGWAS (left panels) and GEMMA (right panels). Results for the mouse and rice datasets are displayed in the top and bottom panels, respectively. The red diagonal line represents the null expectation.

**Figure 5.**
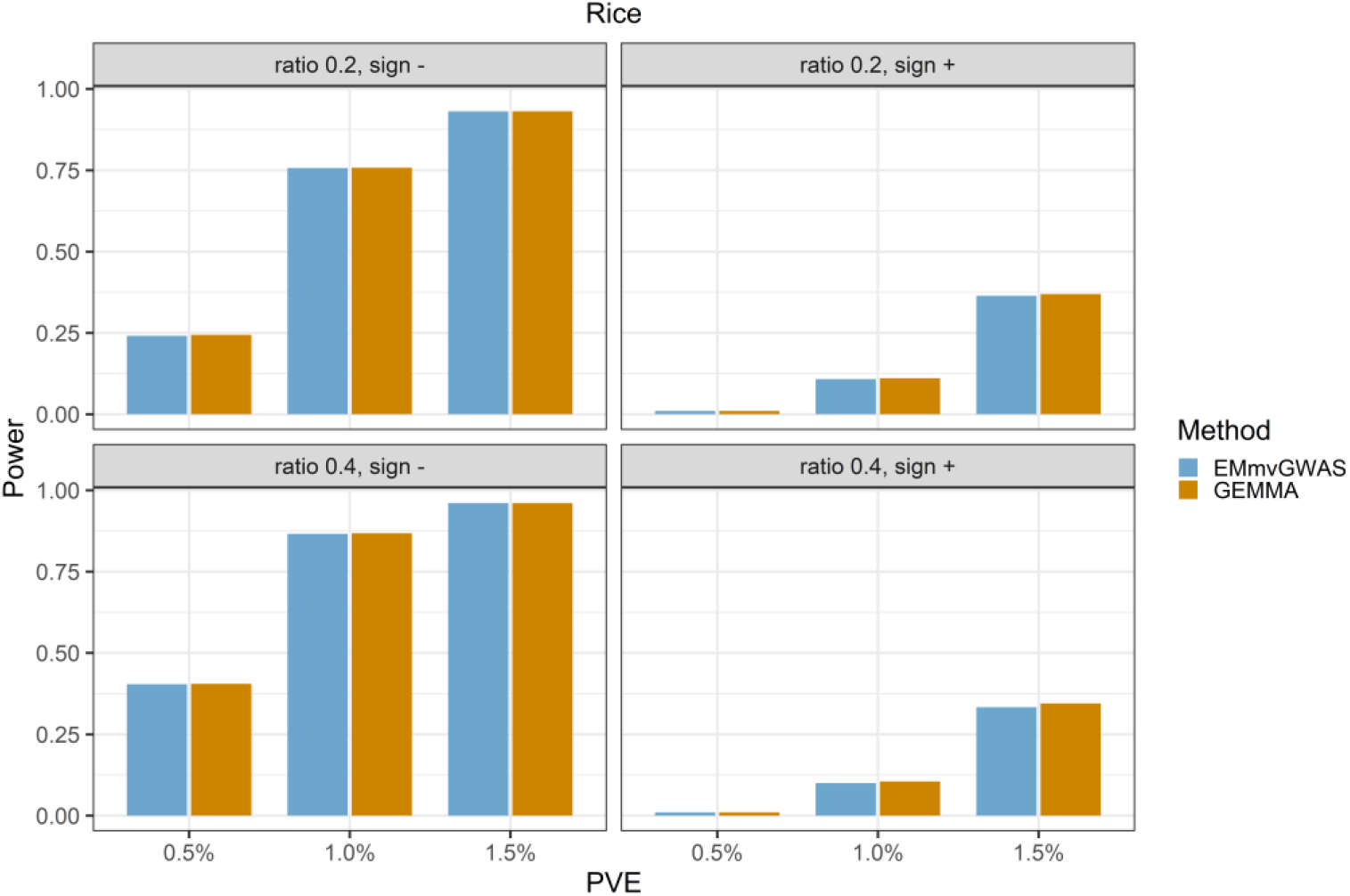
Power comparison between EMmvGWAS and GEMMA under alternative simulations using rice data. Causal SNPs were simulated with varying levels of phenotypic variance explained for the first trait (PVE), and effect sizes on the second trait were set as either 20% or 40% of PVE, with effects in the same (sign +) or opposite (sign –) directions. Power was defined as the proportion of causal SNPs detected at the Bonferroni-corrected significance level.

## 4. Discussion

These results highlight the substantial computational gains offered by EMmvGWAS. While GEMMA scales poorly with the dimensionality of the trait space, the EMmvGWAS framework maintains a nearly constant runtime. This efficiency arises from the closed-form re-estimation of the residual variance, which avoids repeated iterative optimization. The performance advantage is consistent across organisms with diverse sample sizes and genetic architectures, demonstrating the scalability and robustness of EMmvGWAS. The human analysis further demonstrates that the semi-exact framework can efficiently leverage parallel computing, achieving additional reductions in runtime without compromise in accuracy. The efficiency of EMmvGWAS opens the door to comprehensive multivariate GWAS across diverse datasets and trait dimensions.

While exact methods are often expected to produce more significant results, this was not observed in our comparisons. We suspect that GEMMA’s internal calibration procedure reduces the apparent significance of its test statistics, leading to closer agreement with the semi-exact method. QQ plots from null simulations confirmed that EMmvGWAS yields well-calibrated p-values that are comparable to GEMMA, with no evidence of systematic inflation or deflation of the type 1 error. This indicates that EMmvGWAS does not compromise type I error control while offering substantial computational efficiency.

For variance component estimation, we employed the expectation-maximization (EM) algorithm under the pure polygenic (null) model. Despite its simplicity, the EM algorithm provided sufficiently accurate estimates and was computationally efficient in our analyses. In contrast, GEMMA uses a more complex procedure that combines parameter-expanded EM algorithm with Newton–Raphson optimization to accelerate convergence. However, our results demonstrate that the standard EM approach is already fast in practice and does not compromise the accuracy of downstream test statistics. This suggests that, for moderate numbers of traits, the EM-based variance component estimation strikes a favorable balance between simplicity, speed, and reliability.

Our method is implemented in the R programming language to ensure accessibility and ease of use within the statistical genetics community. While R is not optimized for high-performance computing, our implementation still demonstrates competitive efficiency. We encourage future implementation in compiled languages such as C++, which could offer further improvement in speed. Additionally, genome-wide marker scanning is inherently parallelizable, and our package includes built-in support for parallel computing to take advantage of multi-core computing systems. The framework could also be extended to support GPU acceleration, enabling even greater scalability for large-scale multivariate GWAS.

## Declarations

### Ethics approval and consent to participate

This study did not involve the collection of any new human or animal data. All analyses were conducted using previously published datasets with existing ethical approval, including UK Biobank data accessed under an approved application. No additional ethics approval or consent was required for this work.

### Consent for publication

Not applicable

### Availability of data and materials

The data set of 1,495 hybrid rice varieties is available at the Rice Haplotype Map Project database (http://www.ncgr.ac.cn/RiceHap4). The CFW mouse datasets can be available at Dryad Digital Repository (doi:10.5061/dryad.2rs41). UK Biobank (UKB) data can be accessed through the UKB Research Analysis Platform (RAP) following approval via the UKB access system (https://www.ukbiobank.ac.uk).

### Authors’ contributions

SX conceived the proposed method. CT analyzed and interpreted the data and generated the figures in the manuscript. CL and QW provided processed UKB dataset. YC provided the hybrid rice dataset. XW advised on the GEMMA implementation. CT wrote the manuscript. SX supervised the research and edited the manuscript. All authors reviewed the manuscript.

