## Supplementary Table S1 for "An Expectation and Maximization Algorithm for Multivariate Genome-wide Association Studies (EMmvGWAS)"

| Dataset | Original Sample Size | Filtered Sample Size | Original #SNPs | Filtered #SNPs | Traits |
| --- | --- | --- | --- | --- | --- |
| Rice | 1,495 | 1,495 | 201,707 | 181,973 | Yield (YD), Panicle number (PN), Grain number (GN), Seed setting rate (SSR), 1000 grain weight (TGW), Heading date (HD), Plant height (PH), Panicle length (PL), Grain length (GL), Grain width (GW) |
| Mouse | 1,149 | 1,091 | 92,734 | 92,734 | Tibialis anterior weight (TA), Extensor digitorum longus weight (EDL), Body weight at week 1 (bw1), Body weight during prepulse inhibition testing (PPIweight), Tail length, Tibia bone length (tibia), Testis weight, Gastrocnemius weight (gastroc), Plantaris weight, Soleus weight |
| Human | 520,000 | 5,000 | 90M | 2,026,362 | Body mass index (BMI), Weight, Basal metabolic rate, Waist circumference, Hip circumference, Standing height, White blood cell count (WBC), Red blood cell count (RBC), Platelet count, Eosinophil count |

**Table S1. Summary of sample sizes, SNP counts, and phenotypic traits for the rice, mouse, and human datasets.**
